# Improving cross-cultural “mind-reading” with electrical brain stimulation

**DOI:** 10.1101/313478

**Authors:** A. K. Martin, P. Su, M Meinzer

## Abstract

**Background:** A cross-cultural disadvantage exists when inferring the mental state of others, which may be detrimental for individuals acting in an increasingly globalized world. The dorsomedial prefrontal cortex (dmPFC) is a key hub of the social brain involved in ToM. Therefore, we explored whether facilitation of dmPFC function by focal high-definition tDCS can improve cross-cultural mind-reading.

**Method:** 52 (26 F/M) Singaporeans performed the Caucasian version of the Reading the Mind in the Eyes Test (RMET) and received HD-tDCS to either the dmPFC or a control site (right temporoparietal junction,rTPJ) in sham-controlled, double-blinded, crossover studies. Contact with Caucasians was determined for the Singaporean cohort as a potential mediator of RMET performance and HD-tDCS response. 52 Caucasians completed the RMET during sham-tDCS and served as a comparison group.

**Results:** A cross-cultural disadvantage on the RMET was confirmed in the Singaporean cohort and this disadvantage was more pronounced in those participants who had less contact with Caucasians. Importantly, HD-tDCS to the dmPFC improved RMET performance in those with less contact. No effect was identified for rTPJ HD-tDCS or for the age/sex control task demonstrating task and site specificity of the stimulation effects.

**Conclusion:** Electrical stimulation of the dmPFC selectively improves the rate of cross-cultural ToM inference from facial cues, effectively removing cross-cultural disadvantage that was found in individuals with lower cross-cultural exposure.

## INTRODUCTION

In an increasingly globalised world, cross-cultural understanding is of paramount importance. Understanding the underlying beliefs, intentions, and emotions of others requires the attribution of underlying states from observable characteristics, often labelled Theory of Mind (ToM). For example, non-verbal cues are especially rich in information about such underlying states of mind and the eyes and surrounding regions are often thought to hold special prominence in mental state decoding.

The “reading the mind in the eyes test” (RMET) involves inferring the mental state from a facial cue predominantly involving the eye region. Cultural differences in the ability to “read the mind in the eyes” have been identified, with an advantage for own-culture judgements compared to cross-cultural judgements in native Japanese and white Americans (Adams et al., 2010). The Experienced-based perceptual theory (Brigham, Maass, Snyder, & Spaulding, 1982; Chiroro & Valentine, 1995; Malpass & Kravitz, 1969; Valentine, 1991) proposes that a lack of experience with other-race faces leads to a basic perceptual inability to code facial features as accurately as for same-race faces. Although this theory is usually reserved for studies in relation to memory for other-race faces, a lack of experience with Caucasian faces may explain the deficit on the Caucasian version of the RMET in Asian subjects recently arrived into a majority Caucasian country. However, an intra-cultural advantage also exists when presented with more complex mind-reading tasks using verbal vignettes that do not rely on facial features, suggesting that inter-cultural mind-reading is impaired over and above difficulties on lower level perceptual discrimination (Perez-Zapata, Slaughter, & Henry, 2016).

The RMET is associated with increased activity in regions within the ‘social brain’, such as the temporoparietal junctions (TPJ) and the dorsomedial prefrontal cortex (dmPFC; for review see Molenberghs, Johnson, Henry, & Mattingley, 2016; Schurz, Radua, Aichhorn, Richlan, & Perner, 2014). In addition, studies across development from adolescence to adulthood have shown that the dmPFC is engaged during the RMET in adolescence and decreases into early adulthood (Blakemore, 2012; Blakemore, den Ouden, Choudhury, & Frith, 2007; Burnett, Bird, Moll, Frith, & Blakemore, 2009; Moor et al., 2012; Wang, Lee, Sigman, & Dapretto, 2006). This developmental pattern suggests that the dmPFC may play a greater role during the learning phase of acquiring a skill, with other regions taking over later in development or expertise (Johnson, 2011). The right TPJ shows consistent activation across ToM tasks including the RMET (Schurz et al., 2014). However, the TPJ regions do not show the same decline as the mPFC across adolescence into adulthood, providing correlational evidence that its activity is not dependent on familiarity or involved in the learning process (Blakemore, 2012).

In the following study we have used focal, high-definition, transcranial direct current stimulation (HD-tDCS) in a cohort of recently arrived Singaporean students in order to answer the following questions: A) Can we replicate the cross-cultural disadvantage on the RMET, by comparing their performance during placebo (“sham”) HD-tDCS to a carefully matched group of Caucasians students? B) Does performance on the RMET in Singaporeans depend on the amount of contact with Caucasians? and C) can excitatory “anodal” HD-tDCS to the dmPFC remove culturally mediated disadvantage in mind-reading ability in those with less contact?

## METHOD

### Participants

Singaporean students living in Brisbane to complete tertiary study (N=52; 26 M/F) and Caucasian Australian students (N=52; 26 M/F) were stratified equally to either the dmPFC or rTPJ sham-controlled, double-blinded, crossover studies. Singaporeans were selected as English is their first language, thereby removing a potential confound in performance while varying cultural background. Stimulation sessions were counterbalanced so half would receive excitatory “anodal” HD-tDCS in the first session and half in the second session. The sessions were a minimum of three days apart to avoid carryover effects of stimulation (Kuo et al., 2013). All subjects were tDCS-naive, were not currently taking psychoactive medication, and had no diagnosis of neurological or psychiatric disorder. All participants provided written consent prior to inclusion, completed a safety-screening questionnaire and were provided with a nominal fee to compensate their time.

### Transcranial Direct Current Stimulation (tDCS)

HD-tDCS was administered using a one-channel direct current stimulator (DC Stimulator Plus^®^, NeuroConn). Two concentric electrodes were used; a small centre electrode (diameter: 2.5cm) and a ring-shaped return electrode (diameter inner/outer: 9.2/11.5cm). The rTPJ return electrode was smaller (7.5/9.8 cm) to avoid the right ear. HD-tDCS was administered with 1 mA for 20 minutes during the RMET task and only 40 seconds during the sham condition. The latter renders the stimulation ineffective while preserving blinding integrity (Gbadeyan, Steinhauser, McMahon, & Meinzer, 2016). Details of the stimulation parameters and biophysical modelling evidence for focal current delivery with both set-ups are provided in greater detail in previous manuscripts (Martin, Huang, Hunold, & Meinzer, 2017; Martin, Huang, & Meinzer, 2018). Sessions were at least three days apart to avoid carryover effects. Researcher and subject were blinded to the stimulation type. Adverse effects and mood ratings were acquired after each stimulation session.

### Baseline Testing

All participants completed a detailed cognitive battery including the Stroop Test, National Adult Reading Test (NART), Boston Naming Test, phonemic and semantic verbal fluency, and the following tests from CogState^®^ computerized test battery (https://cogstate.com/): International shopping list, Identification test, One-back, Two-back, Set-shifting test, Continuous paired associates learning test, social-emotional cognition test, and the International shopping list - delayed recall. Groups were compared across all cognitive and psychological measures to ensure performance on the RMET or subsequent stimulation effects were not due to underlying differences. Baseline depression and anxiety were assessed using the Hospital Anxiety and Depression Scale (HADS; Zigmond & Snaith, 1983). Social functioning was assessed using the Autism Spectrum Quotient (ASQ; Baron-Cohen, Wheelwright, Skinner, Martin, & Clubley, 2001).

### Reading the Mind in the Eyes Test

The Reading the Mind in the Eyes test (RMET; Baron-Cohen, Wheelwright, Hill, Raste, & Plumb, 2001) is considered a measure of affective Theory of Mind (ToM) functioning, although recently conceptualized as a measure of emotion recognition (Oakley, Brewer, Bird, & Catmur, 2016). Subjects completed the Caucasian version of the RMET as part of a larger battery of social cognitive tests described in detail in a previous study from our group (Martin, Dzafic, Ramdave, & Meinzer, 2017). Subjects completed the same test in session two, although the order of stimuli was randomized to prevent simple memory effects. A set of eyes were presented with four mental attribution words surrounding. Subjects were instructed to choose the mental attribution that was best represented in the eyes. Subjects were then asked to judge the age and sex of the person in the picture, again on a four-point scale (1= young male, 2= young female, 3= old male, 4= old female). Old was considered greater than 50 years of age. We modified the original task as described by (Baron-Cohen, Wheelwright, Hill, et al., 2001) to include a greater variety of female eyes, especially older females. We also amended the response options so that each option only occurred once throughout the test. There was no time limit on responding and the next stimulus was presented as soon as a response was recorded. Participants were able to ask for definitions of words if they did not understand. The age/sex question occurred directly after the mental state attribution question, with both conditions during online (active or sham) stimulation. Accuracy scores were computed for the mental attributions and age and sex judgments (each with a total correct out of 38).

### Contact with Caucasians Questionnaire

Prior to the stimulation, participants completed a questionnaire about their level of contact with Caucasians (Cao, Contreras-Huerta, McFadyen, & Cunnington, 2015). They were required to estimate the percentage of Caucasians in the following social contexts: neighbourhood, university, classmates, and friends. We excluded the workmates question as too few were working at the time of the study. We were only interested in their contact since arriving in Brisbane as their contact with Caucasians in Singapore was very low across the entire cohort (<5%). We took an average score across all social contexts as our Contact with Caucasians measure.

### Statistical Analyses

All analysis was conducted in SPSS version 23 and JASP version 0.8.6. T-tests were used to assess baseline performance on the RMET ToM and Age/Sex control task between Caucasians and Singaporeans. Univariate analysis of variance (ANOVA) was used to identify whether performance was associated with underlying ASQ traits. Pearson’s correlations were performed to assess the relationship between baseline RMET performance and Contact with Caucasians. Partial correlation was used to assess whether differences in the higher-order ToM task were due to ASQ or differences on the lower-order Age/Sex discrimination task. Regression models were computed for both RMET measures with Stimulation Site and Contact with Caucasians as predictors. Significant results were followed up with simple-effects analyses at each Stimulation Site. Partial correlation was used to further scrutinize a specific effect of stimulation on the higher-order ToM task that was not reliant on lower-order face discrimination.

## RESULTS

### Baseline Measurements

The groups were matched for depression and anxiety symptoms, but the South-East Asian group scored higher on the ASQ (see Supplementary **Table 1**). As ASQ was associated with RMET performance with poorer performance in those with higher ASQ scores, (r=-0.20, p=0.04) we controlled for ASQ across analyses where appropriate.

Baseline cognition, ASQ, depression, and anxiety scores were comparable between the South-East Asian groups receiving either dmPFC or rTPJ stimulation, with only a small difference in set-switching ability (p=0.04) (see Supplementary Table 2).

### Reading the Mind in the Eyes

South-East Asians performed worse than the Caucasian cohort on the RMET, t(102)=-2.41, p=0.02, Cohen’s d=-0.47, although this was no longer significant after controlling for ASQ, F(1, 101)=1.61, p=0.21. One South-East Asian outlier was removed from the age/sex judgement analysis as they performed greater than four standard deviations below the South-East Asian mean (inclusion did not alter the results though). Performance on the age/sex judgements was also worse in the South-East Asians, t(101)= -2.42, p=0.02, Cohen’s d=-0.48 and this remained significant after controlling for ASQ, F(1, 100)=4.89, p=0.03.

**Figure 1.**
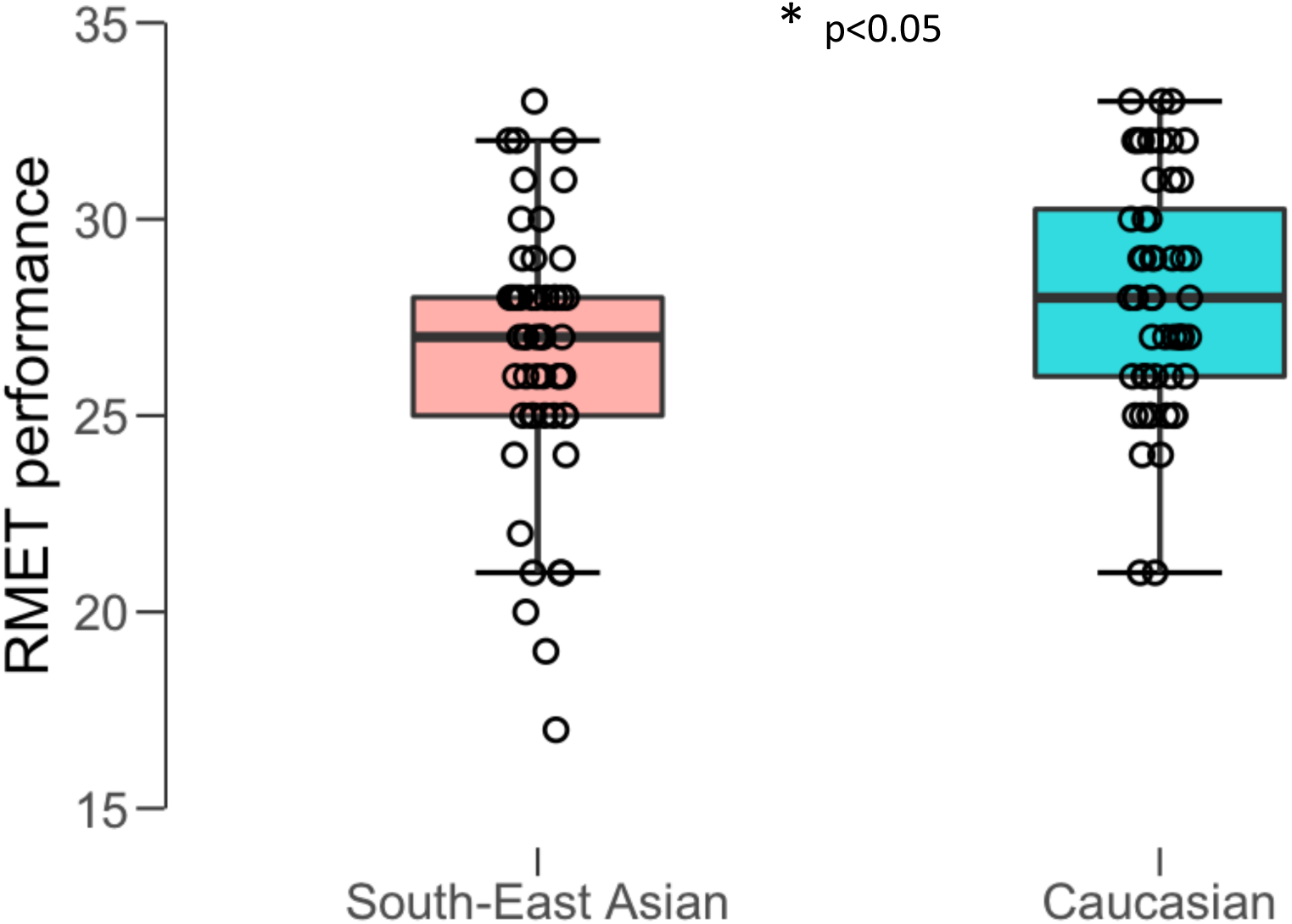
South-East Asian Singaporeans showed a cross-cultural disadvantage on the RMET (26.56 v 28.06) compared to Caucasian Australians.

Within the South-East Asian cohort, performance on the RMET during sham-tDCS correlated with Contact with Caucasians, r=0.35, p=0.008 (see **Figure 2**) and this remained significant after controlling for ASQ, r=0.32, p=0.02. No such relationship was identified for the age and sex judgement task, r=0.16, p=0.28. The significant correlation between RMET and Contact with Caucasians also remained significant after partial correlation controlling for performance on the age/sex judgements, r=0.33, p=0.02. This shows that Contact with Caucasians influences the higher-order affective ToM judgements independent of lower level feature detection or underlying ASQ traits.

**Figure 2.**
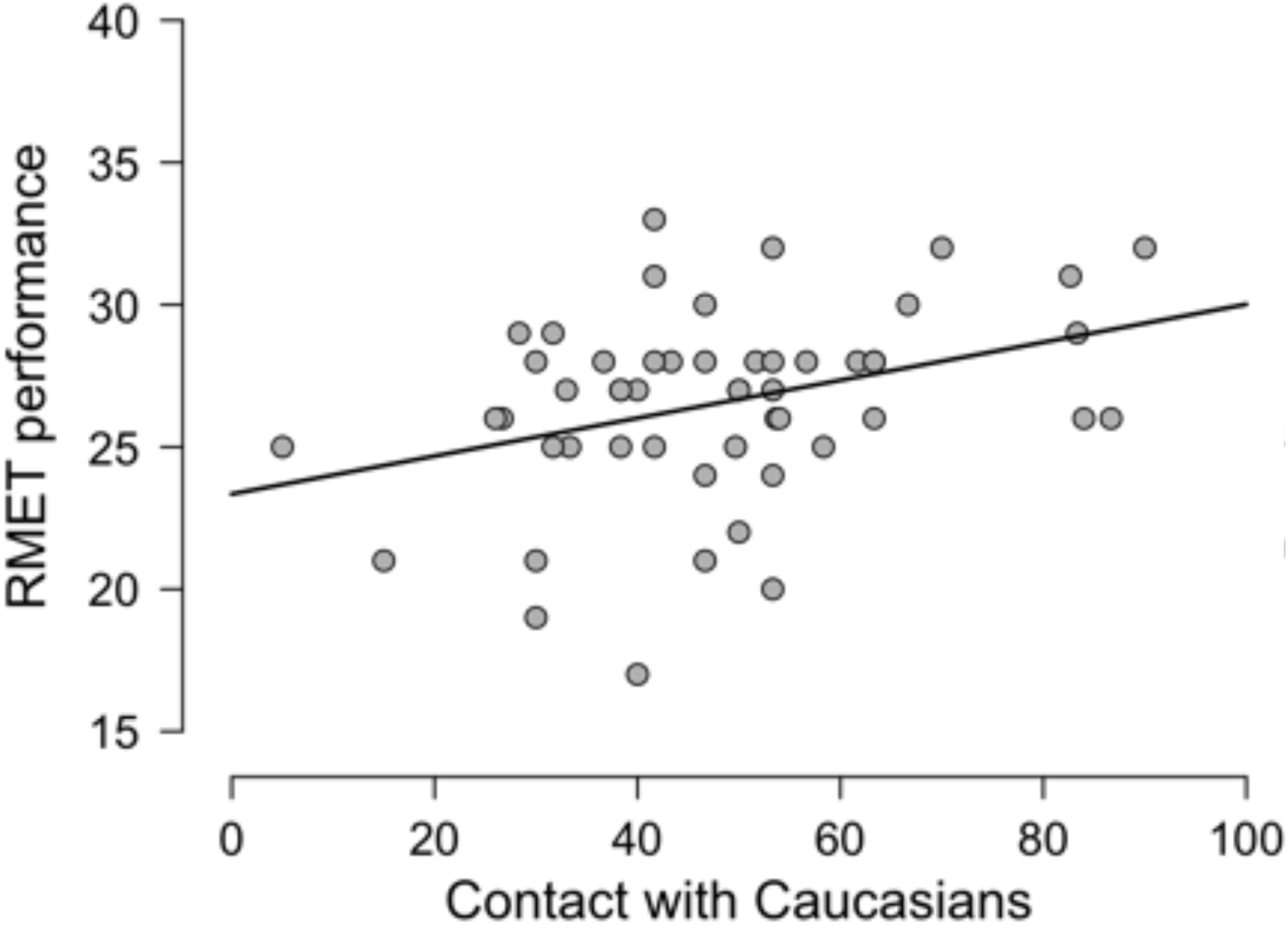
In the South-East Asian group, performance on the RMET during sham HD-tDCS was positively correlated with contact with Caucasians, r=0.35, p=0.008.

A regression model with RMET improvement as the dependent variable and Stimulation Site and Contact with Caucasians was calculated. The model was a suitable fit for the data, F(2,49)=3.78, p=0.03. Only Stimulation Site significantly contributed to the model, t=-2.59, p=0.01. Univariate ANOVAs for both the dmPFC and rTPJ studies revealed that after controlling for Contact with Caucasians, only dmPFC stimulation improved performance on the RMET, F(1,24)=6.28, p=0.02, whereas rTPJ had no effect, F(1,24)=0.27, p=0.61, thereby demonstrating regionally specific stimulation effects.

Furthermore, we inspected the relationship between Stimulation site and Contact with Caucasians independently for the dmPFC and rTPJ. This showed that improvement on the RMET during dmPFC stimulation negatively correlated with Contact with Caucasians, r=-0.412, p=0.036, such that improvement was greater in those with less contact (see Figure 3). There was no relationship between rTPJ stimulation and Contact with Caucasians, r= 0.001, p=0.99. A regression model with Age/Sex judgement improvement as the dependent variable and Stimulation Site and Contact with Caucasians was also calculated but was not significant, F(2,48)=0.99, p=0.38.

**Figure 3.**
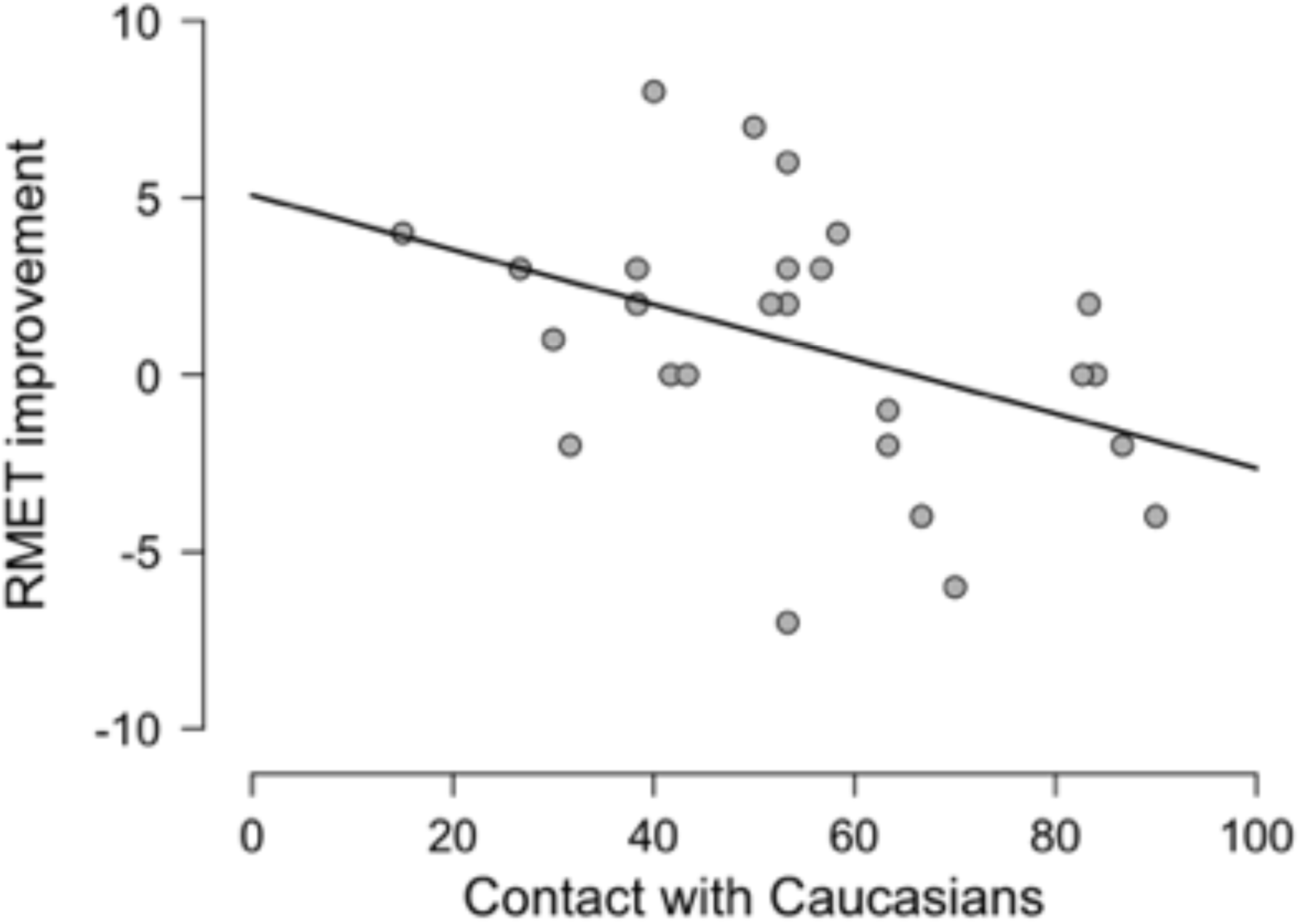
Improvement on the RMET after anodal HD-tDCS to the dmPFC was dependent on Contact with Caucasians. Those with less contact had greater improvement, r=-0.41, p=0.036.

To further clarify whether the improvement on RMET performance, dependent on Contact with Caucasians, was independent of improvement on the lower-order Age/Sex control task, a partial correlation was calculated. The significant correlation between improvement on the RMET and Contact with Caucasians remained significant after partial correlation controlling for improvement on the Age/Sex judgements, r=-0.409, p=0.042. Therefore, dmPFC stimulation improved RMET performance independent of lower-order perceptual improvements.

There were no significant differences between performance on the RMET nor the age/sex attributions between the first and second sessions across the entire sample or in each sample independently (all p>0.6). This shows that potential learning effects over the two HD-tDCS sessions did not influence the results.

### Adverse Effects and Blinding

All participants tolerated the stimulation well with only minor physical sensations. Self-reported mild adverse effects and mood (Brunoni et al., 2011; Folstein & Luria, 1973) were comparable between the stimulation conditions (see Supplementary **Table 3**). Although anodal stimulation resulted in more adverse effects across both rTPJ and dmPFC groups combined (p=0.05), there was no difference between stimulation sites (p=0.67). Subjects guessed the stimulation order at chance level (number of correct guesses: 30/52, p=0.27; there was no difference between stimulation site, p=1.00). This demonstrates that the behavioural effects of stimulation, specific to the stimulation site, were not affected by non-related minor side-effects of the stimulation or the participants’ ability to recognize active stimulation.

### Discussion

This is the first study to identify a beneficial effect of electrical stimulation on cross-cultural mind-reading. We replicated the cross-cultural disadvantage and provided novel evidence that this may be due to higher underlying ASQ traits. We also demonstrate that this disappears with increased cross-cultural contact. Finally, HD-tDCS to the dmPFC can improve RMET performance specifically in those with less contact with Caucasians in the absence of cognitive training. Both task and stimulation site specificity were demonstrated as there was no effect of stimulation on the age and sex discrimination task and no effects on either task during rTPJ stimulation.

The results of the current study replicate previous research demonstrating cross-cultural disadvantage on the RMET (Adams et al., 2010; Perez-Zapata et al., 2016) and provides the first evidence that cross-cultural ToM judgements improve as contact with that cultural group is increased. It suggests that increased cross-cultural exposure may strengthen associations between external physical cues and underlying mental states, improving cross-cultural mind-reading ability. As contact with Caucasians did not predict performance on the lower level age/sex judgements, it suggests that a lower-level perceptual based theory (Brigham et al., 1982; Chiroro & Valentine, 1995; Malpass & Kravitz, 1969; Valentine, 1991) is not sufficient in explaining improvements in cross-cultural mind-reading.

The current study also provides the first evidence that HD-tDCS can improve cross-cultural mind-reading ability, even in the absence of cultural training or exposure. Excitatory stimulation over the dmPFC increased mind-reading performance specifically in those with less recent contact with Caucasians. One possible explanation for this effect is that the dmPFC may be more active in South-East Asians with less contact with Caucasians in the same manner that the dmPFC is more active in adolescents than adults during mentalizing tasks (Blakemore, 2012; Blakemore et al., 2007; Burnett et al., 2009; Moor et al., 2012; Wang et al., 2006). The dmPFC may be required to actively recall paired associations between percepts and their mental labels and that once sufficiently learnt, the dmPFC is no longer required (Johnson, 2011). The dmPFC has also been implicated in representing in-group compared with out-group members (Molenberghs & Morrison, 2014; Morrison, Decety, & Molenberghs, 2012; Telzer, Ichien, & Qu, 2015). Therefore, another possibility is that those with less contact may represent Caucasians more as outgroup members and fail to sufficiently recruit social brain regions such as the dmPFC when inferring mental states. As we have previously identified a role for the dmPFC in the integration of other encoded information into the self (Martin, Dzafic, et al., 2017; Martin et al., 2018), anodal stimulation to the dmPFC in South-East Asian may increase the ability to integrate the mental state of the other into that of the self and improve performance on the RMET.

Importantly, this study only investigated Singaporeans in Australia and future research could assess the consistency of this effect in different cultural groups. As Caucasians also perform worse at mindreading when the eyes are of South-East Asian origin (Adams et al., 2010), it is conceivable that contact with South-East Asians would improve performance in Caucasians and that stimulation of the dmPFC may accelerate this process. As the current study is correlational in regards to performance on the RMET and contact with Caucasians, we are unable to rule out the possibility that greater mindreading ability leads to greater contact with Caucasians, although the percentage of Caucasians in settings such as a neighbourhood, university, and classroom are not easily manipulated.

In sum, cross-cultural disadvantage on the RMET test may disappear after sufficient contact with the cultural group in question. This learning process may be mediated by the dmPFC and anodal HD-tDCS to this region may accelerate the process, e.g., by enhancing the outcomes of cross-cultural sensitivity training (Pruegger & Rogers, 1994). With inter-cultural migration an increasing part of a globalized world, facilitating cross-cultural understanding holds great promise for harmonious and cohesive societies. This can be achieved by understanding the fundamental human process of ToM, how this differs across and between cultures, and providing cognitive and neuroscientific evidence for ToM improvement.

